# Cooperation between CRISPR-Cas types enables adaptation in an RNA-targeting system

**DOI:** 10.1101/2020.02.20.957498

**Authors:** Ville Hoikkala, Janne Ravantti, César Díez-Villaseñor, Marja Tiirola, Rachel A. Conrad, Mark J. McBride, Lotta-Riina Sundberg

## Abstract

CRISPR-Cas immune systems adapt to new threats by acquiring spacers from invading nucleic acids such as phage genomes. However, some CRISPR-Cas loci lack genes necessary for spacer acquisition, despite apparent variation in spacer content between strains. It has been suggested that such loci may use acquisition machinery from co-occurring CRISPR-Cas systems. Here, using a lytic dsDNA phage, we observe spacer acquisition in the native host *Flavobacterium columnare* that carries an acquisition-deficient subtype VI-B locus and a complete subtype II-C locus. We characterize acquisition events in both loci and show that the RNA-targeting VI-B locus acquires spacers *in trans* using acquisition machinery from the DNA-targeting II-C locus. Our observations reinforce the concept of modularity in CRISPR-Cas systems and raise further questions regarding plasticity of adaptation modules.

---

CRISPR (Clustered Regularly Interspaced Short Palindromic Repeats) arrays consist of different instances of a repeated sequence separated by unrepeated spacer sequences of a regular size (*1*). Together with Cas (CRISPR associated) proteins (*2*), they constitute CRISPR-Cas systems that protect bacteria and archaea against infections by bacteriophages (phages) (*3*). Immunity operates under three main phases: (i) adaptation, (ii) expression and (iii) interference. In the adaptation (or acquisition) phase (i), a fragment from the phage genome (protospacer (*4*)), next to a protospacer adjacent motif (PAM) (*5*), is inserted into a CRISPR array as a spacer. New spacers are acquired at the end of the array next to an AT-rich region called the leader (*2*–*4*, *6*). After acquisition, processed transcripts from the array (ii) guide the recognition of sequences complementary to the spacer in the invading genome, which triggers the activation of nucleases (iii) (*7*–*9*).

CRISPR-Cas systems are divided into six types and several subtypes depending on the *cas* gene composition of their interference modules (*10*). Adaptation, by contrast, is almost universally (*10*) mediated by a spacer acquisition complex consisting of Cas1 and Cas2. This complex interacts with an array’s leader-repeat junction to insert new spacers (*11*). As repeats and leaders vary in sequence and in length, their respective adaptation proteins co-evolve accordingly (*12*–*16*). Therefore, individual CRISPR-Cas loci co-occurring a genome have distinct versions of Cas1 and Cas2. Interestingly, some CRISPR-Cas loci lack adaptation modules but still appear to have variable spacer content in different bacterial isolates (*13*, *17*, *18*). It has been suggested that such loci may use acquisition machinery from other intragenomic CRISPR-Cas loci (*17*, *19*–*23*). However, a major restraint for this *in trans* adaptation model are the aforementioned compatibility requirements regarding leader and repeat sequences. *In trans* acquisition between CRISPR-Cas types is therefore not trivial, and to our knowledge experimental evidence to support this model has not been presented.

The recently discovered type VI CRISPR-Cas systems (*19*, *21*, *24*) often lack *cas1/2*, especially in subtype VI-B, and have thus been proposed to acquire spacers *in trans* (*19*, *21*–*23*). However, as type VI loci exclusively target RNA, obtaining spacers from dsDNA (the *modus operandi* for most acquisition complexes) is not optimal, as only half of potential spacers may functionally target the mRNA strand. In addition, as acquisition complexes cannot directly use RNA as substrate for spacer insertion, immunity against RNA phages cannot be generated. Some type VI-A loci may circumvent these problems through a reverse-transcriptase fused to Cas1 (*25*), but this is not common for most type VI loci (*25*). Adaptation in type VI has not been experimentally demonstrated and it is unknown if new spacers are acquired through the *in trans* adaptation model.

The genome of the fish pathogen *Flavobacterium columnare* has a Cas1/2-deficient VI-B subtype (*cas13b* and an array), accompanied by a complete II-C subtype (*cas1*, *cas2, cas9* and an array) (Fig. 1). The subtype II-C locus uses Cas9 for dsDNA interference, while the subtype VI-B likely promotes cellular dormancy (*26*) through the non-specific RNAse activity of Cas13b (*24*). Previously, we showed that natural isolates of *F. columnare* vary in their phage-targeting spacer content in both loci, driving phage evolution (*18*). Only the phage mRNA strand was targeted in VI-B viral protospacers (n=15), while this was true for only half of the II-C protospacers (*18*). It is unclear if the bias for mRNA-targeting type VI spacers stems from selective acquisition or from positive selection of successful interference. To further characterize spacer acquisition in these loci, we co-cultured *F. columnare* with a lytic dsDNA phage in controlled laboratory conditions. Our results reveal details of subtype VI-B and II-C adaptation in a native host and show the first experimental evidence of *in trans* adaptation between CRISPR-Cas types, despite strong differences in leader and repeat sequences (Fig. 1).

**Figure 1.**
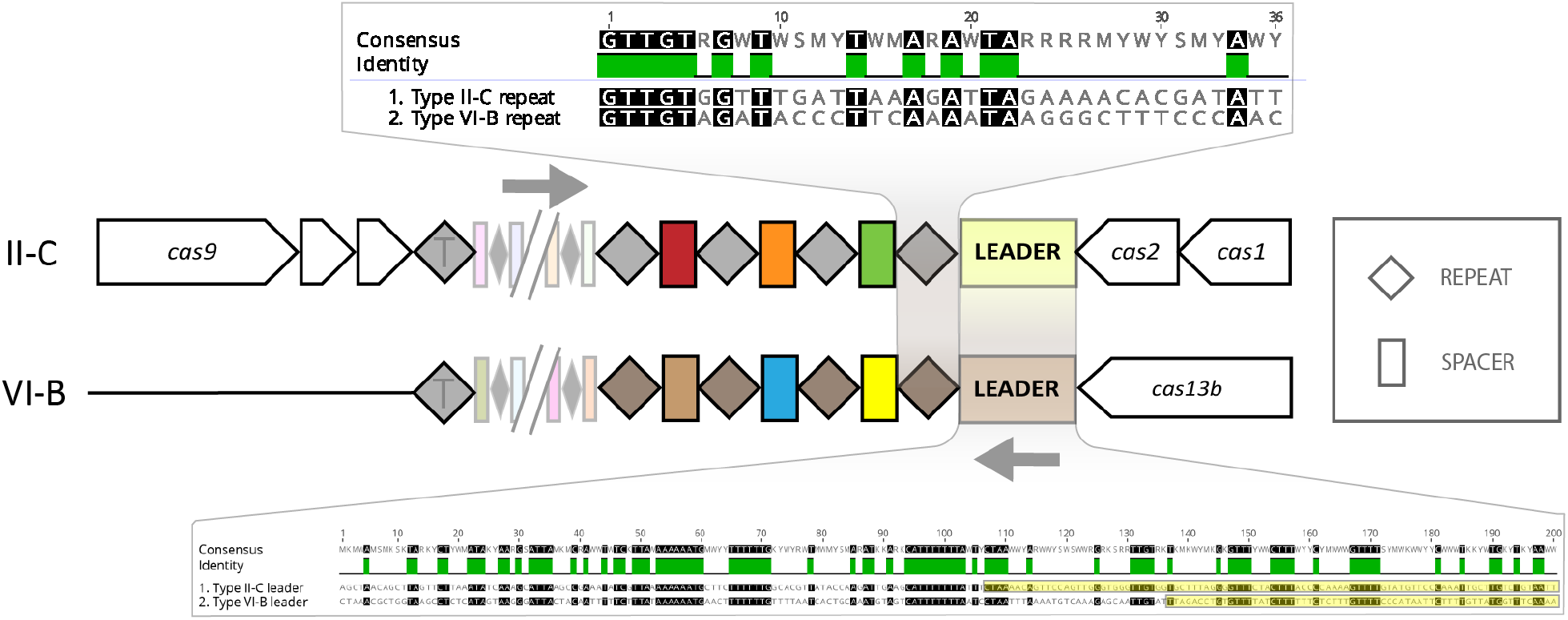
Comparison of the arrangements of II-C and VI-B CRISPR-Cas loci in *F. columnare.* Colored boxes represent spacers and diamonds represent repeats (T denotes the terminal repeat). Repeats and leaders are aligned with no gaps and presented in 5’ to 3’ direction. Yellow annotation in the leaders shows an open reading frame. Annotations not to scale.

## Phage-induced acquisition in both loci

To study the adaptation phase of both subtype II-C and VI-B loci in a controlled environment, we cultured *F. columnare* strain B185 (*27*, *28*) with its virulent dsDNA phage FCL2 (*28*) in liquid medium. We also included treatments with UV-radiated (*29*) FCL2 and a mixture of radiated and non-radiated phage. After a week, we plated the liquid cultures to screen for expanded CRISPR arrays in individual colonies. Only bacteria that were exposed to the phage mixture had expanded arrays, although most of these colonies were still ancestral in their spacer content. We did not observe spacer acquisition in bacteria that were exposed to either phage type alone or in the bacteria-only treatment (Table S1 and S2).

Next, we focused on CRISPR adaptation on the bacterial population scale by amplifying the variable ends of both arrays directly from the liquid cultures. Similar to the colony screen, we saw spacer acquisition in the phage mixture treatment, but also in the non-radiated phage treatment. Within each replicate (identified by letters a to e), the relative efficiency of spacer acquisition was similar between the two arrays (Fig. S1).

To identify the diversity and origin of the new CRISPR spacers in the bacterial population, we deep sequenced the population-scale variable ends of the arrays in the non-radiated phage treatments. We obtained data on new subtype II-C spacers from two replicates (b and e), and subtype VI-B spacers from three replicates (b, d and e). Most II-C spacers targeted the phage genome, with a minority targeting the bacterial genome (Fig. 2). By contrast, VI-B spacers targeted the bacterial and phage genomes evenly. The different targeting-preferences between the loci are likely explained by negative selection of self-DNA cleavage (*30*), which is induced by Cas9 but not by Cas13b.

**Figure 2.**
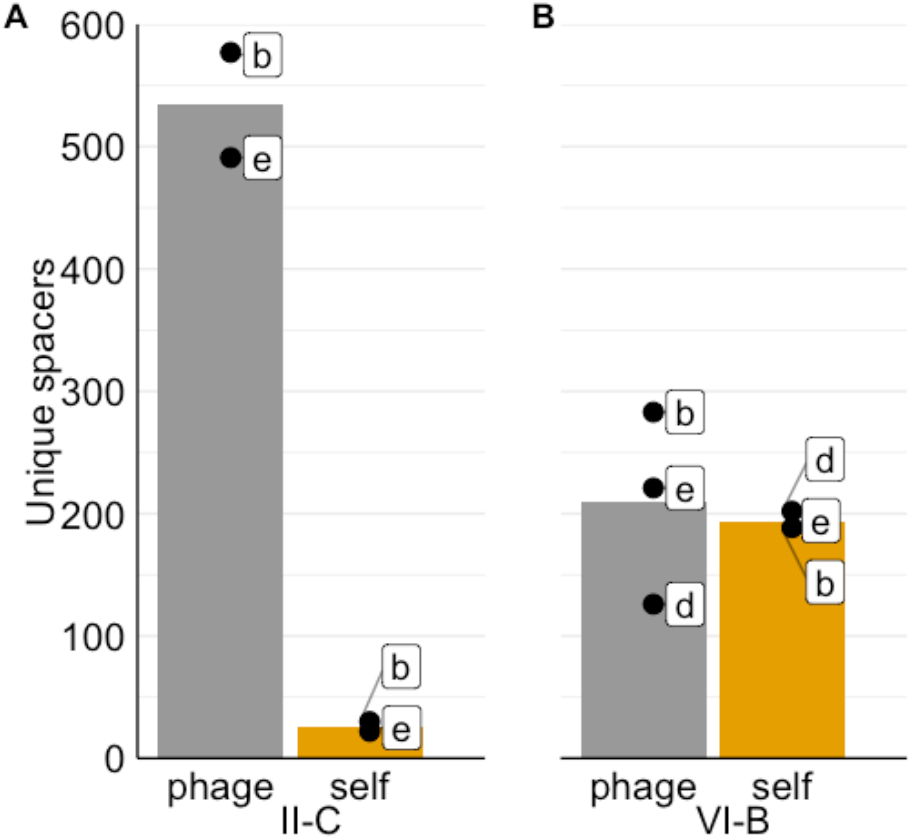
Population-level analysis of origin of unique spacers. Number of new unique subtype II-C (A) or VI-B (B) spacers targeting the phage or bacterial genome. Dots show the exact counts of spacers in a replicate and bars show their respective mean.

To calculate the proportion of identical new spacers between the loci, we compared pooled phage-targeting spacers from both loci. Remarkably, 58% of subtype VI-B spacers were also found in the larger subtype II-C spacer pool. For statistical comparison, we simulated spacer acquisition across the phage genome by sampling random positions freely or next to predicted PAM sites (see below). We calculated the proportion of shared spacers between the simulated spacers and the observed subtype II-C spacers: 6.3% (SD 1.2%) of the random spacers and 29.6% (SD 1.85%) of the PAM-adjacent spacers had a match in the subtype II-C spacer set (Fig. S2). The observed similarity for subtype VI-B spacers is significantly higher than for either simulated pool (P < 10^−10^, one-tailed test), which suggests that new spacers are sampled from a biased protospacer pool that is common to both loci.

## The loci share PAM sequences

To further characterize the protospacers, we investigated PAM sequences for both loci. In the subtype II-C locus, the previously reported pattern (*18*, *31*) NNNNNTAAA was observed downstream of most (~63%) phage-targeting spacers (Fig. S3 and S4). In the few self-targeting subtype II-C spacers, the canonical PAM was almost always either absent or had extended or shortened N-regions, suggesting that both the sequence TAAA and the length of the N-region in the PAM are crucial for subtype II-C interference in *F. columnare* (Fig. S4). The subtype VI-B protospacers had the complementary PAM sequence upstream of the protospacers (TTTANNNNN). This switch in the PAM is explained by opposed transcription directions of the loci (Fig. S5), as the PAM ends of the spacers are equally oriented towards the leader in both arrays. PAM conservation was slightly lower in the subtype VI-B protospacers than in the subtype II-C protospacers (~50%). Protospacer flanking sites (PFS), previously found downstream of type VI protospacers(*21*, *24*), were not detected in our samples.

## Spacer target positions are biased

Despite the relatively even distribution of PAM sequences, spacer targets were not uniformly prevalent along the bacterial or phage genomes. On the bacterial genome of *F. columnare*, spacers from both loci congregated on two hotspots (Fig. 3c-d): one was centered on the subtype II-C CRISPR-Cas locus, while the other was in the predicted origin of replication (oriC). A two-hotspot pattern with similar targets was previously found in a subtype I-E system of *E. coli* and was attributed to high concentration of free DNA-ends in these regions (*32*).

**Figure 3.**
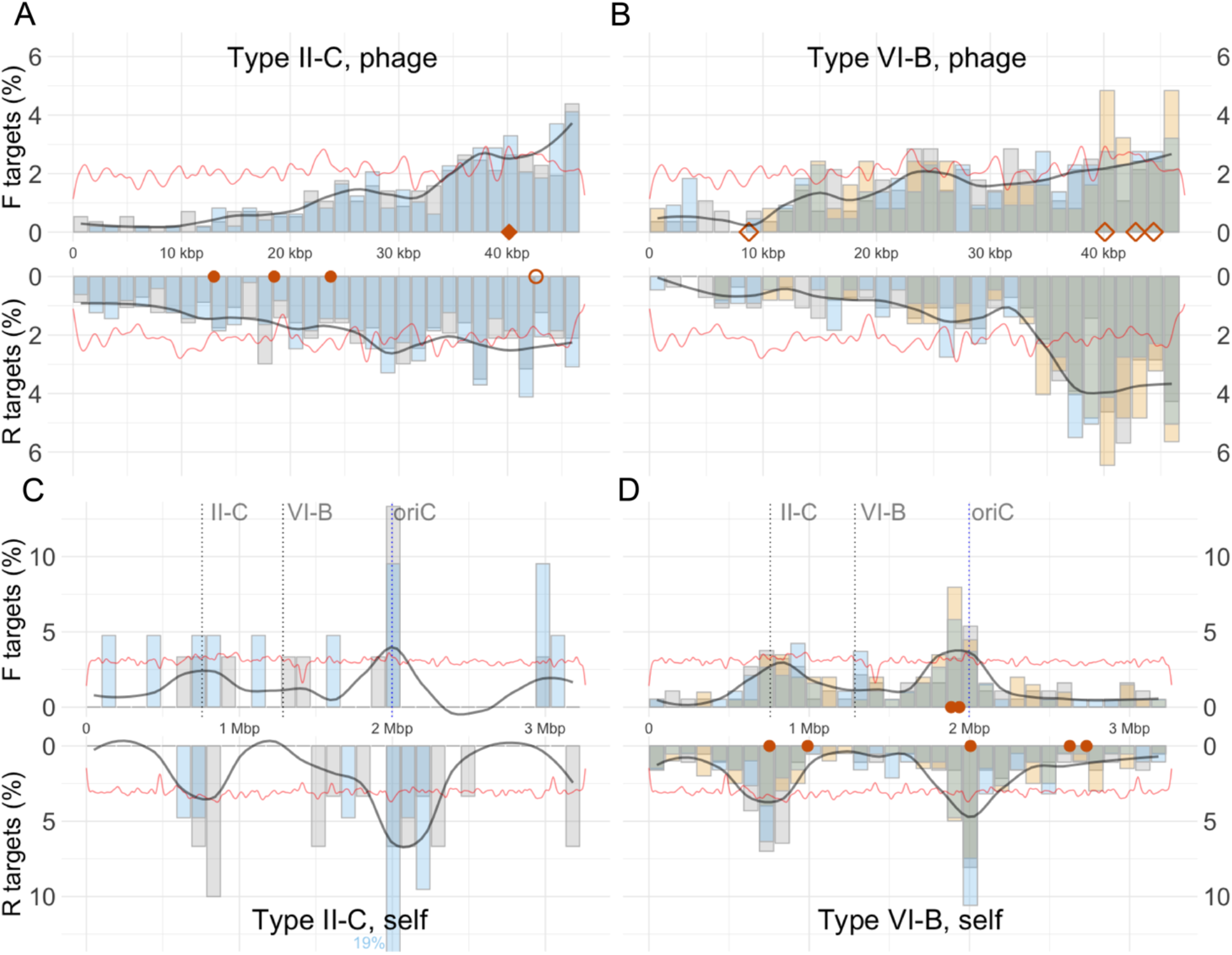
Distribution of unique spacers from subtype II-C and VI-B on phage (a,b) and host (c,d) genomes. The genomes are divided into bins and the targeting percentage of each bin per replicate is shown by overlapping bars (two replicates in subtype II-C, three replicates in subtype VI-B). The black line indicates smoothed average of targeting across replicates. The red line is the relative frequency of the putative PAM sequence. Red markers are spacers that pre-exist in strain B185: diamonds target mRNA and circles do not. Non-filled markers are spacers with 1-3 bp mismatches with their target sequence.

On the phage genome, spacer targets were biased towards the terminal end (Fig. 3a-b). Similar patterns were previously observed in natural isolates of *F. columnare* (*18*) and in the subtype II-A CRISPR-Cas locus of *Staphylococcus aureus* (*33*). The two strands (F and R) of the dsDNA phage genome were targeted unevenly, with stronger bias at the end of the genome (downstream of ~30 kbp). Interestingly, the bias was mirrored between the two loci: as the subtype II-C spacers increased targeting on one strand, the subtype VI-B spacers did so on the other. As suggested above, this can be explained by a common adaptation-bias that is transformed into a swapped strand-bias in interference due to opposing transcription directions of the arrays.

## Most subtype VI-B spacers are nonfunctional

We then investigated if the new spacers reflect the different target-requirements of the loci (DNA vs RNA). The subtype II-C spacers can potentially target either DNA strand, whereas the subtype VI-B spacers must target the coding strand (and thus mRNA) to be functional. On the phage genome, spacers on strand F (Fig. 3a-b) are mRNA-targeting, as all open reading frames (ORFs) are coded by this strand. We would therefore expect the mirrored targeting-patterns (Fig. 3a-b) between the loci to favor mRNA-targeting for the subtype VI-B locus. However, the subtype VI-B strand-bias peak (> 30 kbp) is towards the R-strand and therefore against mRNA-targeting, suggesting that other factors contribute to this pattern.

For more detailed analysis, we examined the capability of the spacers to bind mRNA by assessing their complementarity to predicted ORFs from both target genomes. Spacers from all replicates within a locus were pooled together for an overall estimate. For statistical context, we calculated the probabilities of the different mRNA-targeting proportions with a binomial distribution using sample sizes matching those of the pooled spacers and an equal probability of targeting each strand. We excluded intergenic spacers from this analysis.

Surprisingly, the proportion of functional phage-targeting subtype VI-B spacers was slightly less than half (0.46). The chance of randomly obtaining a proportion at least this low was 7.4% (P=0.074, one-tailed test), suggesting a weak bias towards nonfunctional spacers. Functional self-targeting VI-B spacers, however, were rarer and significantly below expected (P < 10^−9^, one-tailed test) (Fig. 4b).

**Figure 4.**
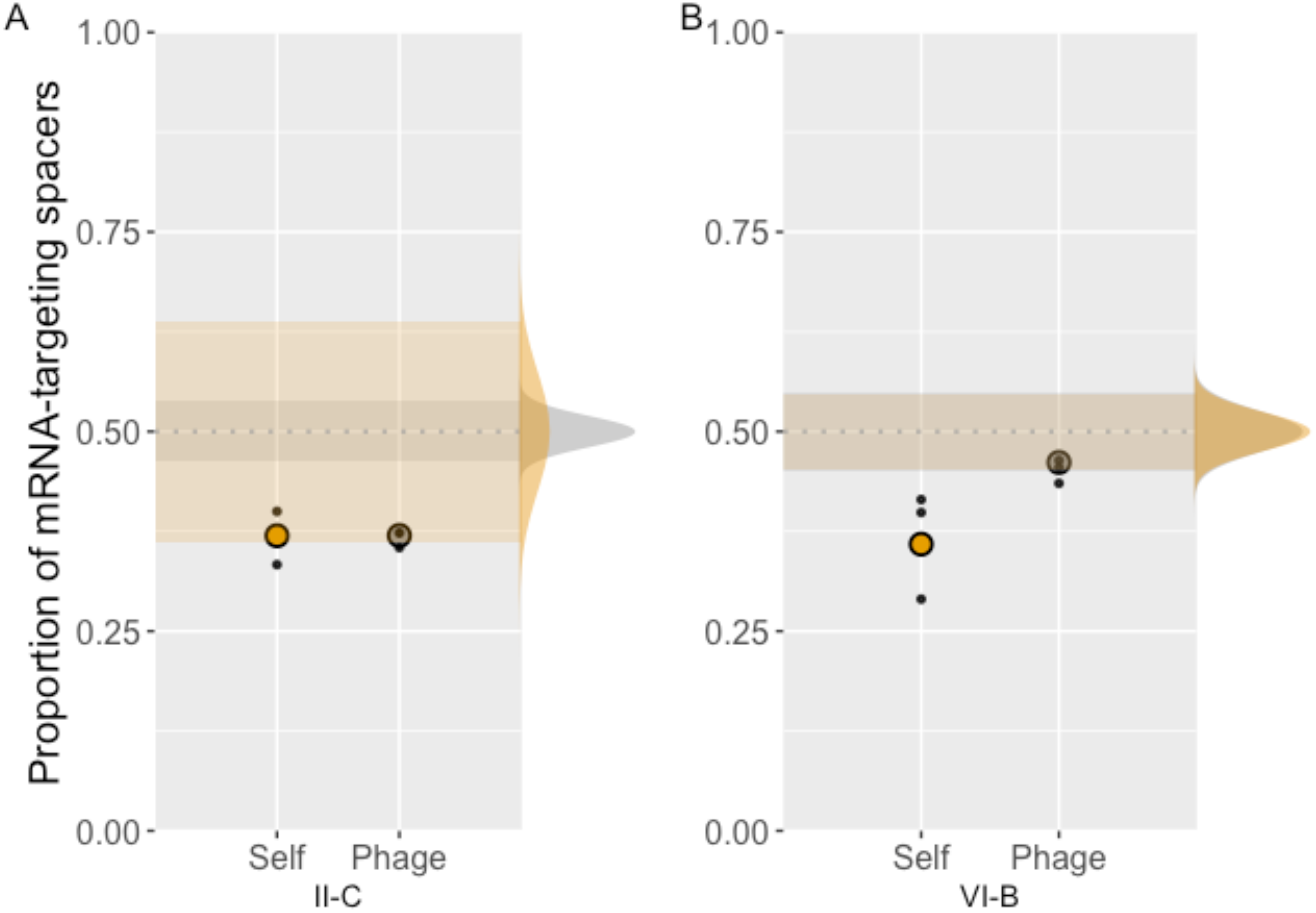
The proportions of ORF targeting spacers that are complementary to mRNA. A) II-C spacers. B) VI-B spacers. Dots are replicates and circles are pooled replicates. Distributions on the right show the spread of expected values adjusted for the size of each spacer pool (yellow = self, grey = phage). 95% confidence intervals of these distributions are shaded on the plot.

Subtype II-C proportions were similar for both target genomes (0.37). Given their vastly different sample sizes, the proportion of mRNA-targeting spacers on the phage genome was significantly below expected (P<10^−10^, one-tailed test), while on the self-genome the proportion was only weakly biased (P=0.052, one-tailed test) (Fig. 4a).

The counter-selection of functional subtype VI-B self-targeting spacers is expected due to their dormancy-inducing effects (*26*) that likely transpire even in the absence of phage. This is also in agreement with the lack of functional self-targeting subtype VI-B spacers in the wild-type array of B185 (n = 7, Fig. 3d). However, this counter-selection is much weaker than for self-targeting subtype II-C spacers (Fig. 2), which argues for a lesser effect of subtype VI-B autoimmunity in this experimental setting.

The avoidance of targeting phage transcripts in subtype II-C spacers is unexpected. A potential explanation would be an uneven distribution of PAM sequences on the two strands, but this is not the case in either genome. Another possible reason for the bias is the inability of RNA-bound Cas9 to degrade DNA: competition between RNA and DNA targets for Cas9 association would impact subtype II-C immunity negatively. The rarity of self-mRNA-targeting II-C spacers could be explained by *in trans* interference: if subtype II-C transcripts were compatible with Cas13b, self-mRNA-targeting spacers would likely be counter-selected.

The reasons for unexpectedly low levels of mRNA-targeting in both loci remain speculative, as targeting-biases shown here are likely affected by multiple factors. However, our results suggest that the complete saturation of functional phage-targeting subtype VI-B spacers in natural isolates (n=19) (*18*) (including the pre-existing four spacers in the wild-type array of B185, Fig. 3a-b) are likely the result of positive selection during interference rather than biased sampling of functional spacers during adaptation. This saturation is not shown in the current study likely due to the short time span of the experiment.

## The VI-B array uses II-C Cas1 *in trans*

Identical acquisition efficiencies, similar protospacer localization patterns and shared PAM sequences supported the hypothesis that the *F. columnare* subtype II-C and VI-B CRISPR-Cas loci share acquisition machinery. To verify this, we deleted *cas1* from the subtype II-C locus to create strain Δ*cas1* and performed a prolonged adaptation experiment with ten phage-exposed and two bacteria-only replicates. As a control, we used a “reversion wild-type” strain (rev-wt) that resulted from an alternative outcome of the mutation process (excision of the chromosomally integrated deletion plasmid can result in either deletion of the target gene or restoration of the wild type sequence). We did not observe population-level spacer acquisition in either array in the Δ*cas1* cultures (Fig. S6). Of all 12 rev-wt cultures, only two phage-exposed replicates survived, and both showed strong spacer acquisition in both arrays (Fig. S6). Despite the unexpected extinction of most rev-wt cultures, these results confirm the dependency of the VI-B system on the II-C acquisition machinery.

## Shared motifs across loci and species

Next, we looked for clues on how the subtype II-C adaptation complex may interact with the subtype VI-B array. When inserting new spacers to the CRISPR array, the Cas1-2 complex depends on conserved sequences in leaders and repeats. In the *Streptococcus thermophilus* subtype II-A CRISPR-Cas locus, the first ten nucleotides on both sides of the leader-repeat junction are essential for spacer integration (*34*) as are the first 41 to 43 nucleotides of the leader in the *E. coli* subtype I-E locus (*11*, *35*). Recently, subtype I-D leaders were also shown to contain important acquisition motives that were more repeat-distal (*36*). Given the lack of similarity in the repeat-leader junctions of *F. columnare* subtype II-C and VI-B loci (Fig. 1), we hypothesized that other regions may show conservation and be important for spacer insertion. We aligned the repeats and leaders of nine species that carry these loci (lacking *cas1* and *cas2* in subtype VI-B) (Fig. S7 and S8). The subtype VI-B repeats were similar in length and sequence, whereas subtype II-C repeats had variation in both. Contrary to expectations, the leader-distal end was more conserved across species in both loci. *F. columnare* was the only species with identical repeat lengths for both loci: other species’ subtype VI-B repeats were paired with subtype II-C repeats that were roughly ten nucleotides longer. As the generation of new repeats is based on precise ruler-mechanisms in the acquisition complex (*35*, *37*, *38*), it remains unknown how these species generated their subtype VI-B array and if the acquisition complex can be tweaked to accommodate two different lengths.

The leaders were more divergent than the repeats. For meaningful alignment, we first separated the species into clusters on the basis of Cas1 similarity, as Cas1 is expected to co-evolve with its respective leader sequence (*12*, *16*). Again, *F. columnare* was an outlier, having the most diverged Cas1 (Fig. S8). Alignments of leaders of both types revealed mostly repeat-distal motifs in the leaders with overall similarities ranging from 38.7% to 58.6%. In *F. columnare* and most other species, the clearest motives were poly-A and poly-T regions at ~50 bp and ~100 bp, respectively (Fig. S8). While speculative, these findings highlight possible binding or signaling sites for spacer integration in both loci using the same acquisition machinery.

## *F. columnare* CRISPR-Cas model

Based on our observations and previous biochemical studies, we propose a model for *F. columnare* spacer acquisition and interference. New spacers are bound to the Cas1-2 acquisition complex in a specific conformation relative to the PAM (*39*, *40*). This pre-spacer is inserted into the array (*38*, *40*–*42*), with the PAM end of the spacer facing the leader (*5*, *43*, *44*). Leader recognition is probably mediated by shared motifs in the subtype II-C and VI-B leaders and repeats that are distal from the leader-repeat junctions. Acquisition results in a downstream interference-PAM for subtype II-C spacers, as is generally observed in type II systems (*45*). Since the arrays are transcribed in opposing directions in respect to the variable end of the array (Fig. S5), the PAM location is switched for subtype VI-B spacers during interference. The same acquisition pattern can therefore lead to two complementary interference patterns, dictated by the array in which the spacer is inserted (Fig. 6). Mirroring of the interference patterns is visible on the phage spacer target maps (Fig. 3).

**Figure 6.**
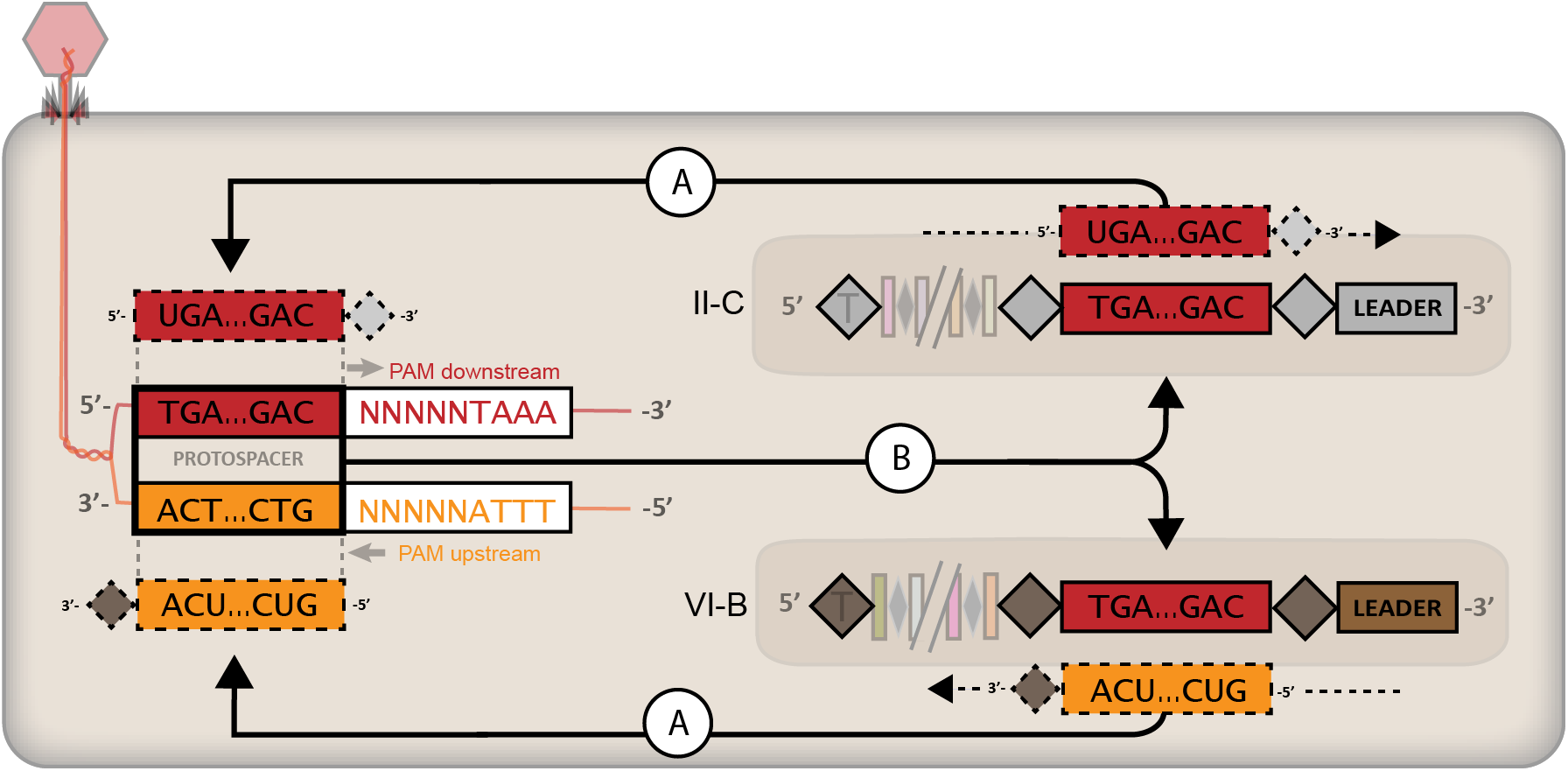
A model for subtype II-C and VI-B acquisition and targeting. A) Matching a spacer to the target genome reveals a downstream PAM for the subtype II-C spacers and an upstream PAM for the subtype VI-B spacers. B) Despite targeting different strands, the original pre-spacer that is inserted may be identical between the arrays. RNA products and transcription are shown with dashed lines.

## Conclusions

We demonstrate that a native subtype VI-B CRISPR-Cas locus acquires spacers from a virulent dsDNA phage and from the host genome. Despite formidable differences in repeat and leader sequences, shared characteristics of the newly acquired spacers with the co-adapting subtype II-C locus supported the model of *in trans* spacer acquisition in the subtype VI-B locus and was confirmed by *cas1* deletion. This arrangement provides the subtype VI-B array with an abundance of nonfunctional spacers and unnecessary carry-over of PAM sequences but spares it the cost of maintaining an adaptation module. The capability to utilize other adaptation modules enforces the concept of modularity in CRISPR-Cas systems that has so far been experimentally shown on the interference level of the immune response (*46*–*48*). We complement these findings with the first case of demonstrated inter-type CRISPR adaptation and also highlight *F. columnare* as one of the few species (*49*) that can be natively induced to acquire spacers from lytic phages in laboratory conditions. We expect future studies to reveal the extent of modularity in type VI systems and the biochemical basis for the plasticity of the acquisition machinery when interacting with different leader and repeat sequences.

## Acknowledgments

We thank Sylvain Moineau for comments on the manuscript, Petri Papponen and Elina Laanto for conducting the RNA sequencing, and Elina Virtanen for Ion Torrent preparation.

## Funding

This work received funding from Kone Foundation, Jane and Aatos Erkko Foundation and the Academy of Finland (grant #314939). This work resulted from the BONUS Flavophage project supported by BONUS (Art 185), funded jointly by the European Union (EU) and the Academy of Finland, and was financially supported, in part, by United States Department of Agriculture-ARS CRIS project 5090-31320-004-00D, cooperative agreement #5090-31320-004-03S to MJM and RAC. The views contained in this document are those of the authors and should not be interpreted as necessarily representing the official policies, either expressed or implied, of the U.S. Government. Mention of trade name, proprietary product, or specific equipment does not constitute a guarantee or warranty by the USDA and does not imply its approval to the exclusion of other products that may be suitable. This manuscript is submitted for publication with the understanding that the United States Government is authorized to reproduce and distribute reprints for governmental purposes. The USDA is an equal opportunity employer.

## Author contributions

Study and experimental design by VH, JR and L-RS. Cas1-mutant planned and executed by RAC and MJM and finalized by VH and CDV. Ion Torrent sequencing planned and assisted by MT. Other laboratory work by VH. Data and statistical analysis by VH with assistance from CDV, JR and LR-S. VH wrote the manuscript with edits and comments from all authors.

## Competing interests

Authors declare no competing interests.

## Data and materials availability

The genomes of *F. columnare* strain B185 (NZ_CP010992.1) and phage FCL2 (NC_027125.1) are available in Genbank. Other data and custom code are not available for the pre-print.

## Supplementary Materials

Figures S1-S8 Tables S1-S3

## Materials and methods

### Bacterial cultures and sampling

Two strains were used to test for CRISPR spacer acquisition: *Flavobacterium columnare* B185 and B245 (*28*). Strain B245 was used in the *cas1*-deletion mutant experiments (see below) and strain B185 was used as the host in the main spacer acquisition study. In the main study, strain B185 was revived from a freezer stock in an overnight SHIEH medium culture (*50*). Five replicates of 5 ml cultures were then inoculated in 0.1X SHIEH medium with the overnight culture to produce an initial concentration of 10^4^ cfu/ml. Phage FCL2 (*28*) was added to the samples at MOI 1. For the phage UV-treatment (*51*), phage FCL2 was exposed to UV light for five minutes on a Petri dish (5000 µJ, UV Stratalinker 1800, Stratagene), lowering infectivity by roughly two orders of magnitude (data not shown). In the bacteria + phage + uv-phage treatments, phages were mixed with an equal volume of UV-treated phages from the same stock and dilution. Five bacteria-only and bacteria + UV-phage cultures were established as controls. The samples were grown at 120 rpm at room temperature for five days, after which they were transferred (1:100) to 5 ml of 1x SHIEH. After two days (day seven total time), cell debris had congregated on the bottom of the tubes. We sampled 1 ml of the clear phase for living planktonic cells and extracted DNA using DNeasy Blood & Tissue Kit (Qiagen). The cultures were also plated on SHIEH-agar plates with dilutions for cfu/ml -estimation and colony-PCR.

### Colony-PCR

The variable ends of type II-C and type VI-B were screened from all conditions (total of 281 colonies) at day 7 (Table S2). The variable ends of both loci were amplified with the primers C1_F + C1_R and C2_F + C2_R (Data S3). Colonies with expanded arrays went through two serial platings and the variable ends were re-checked. Colonies that still showed expanded arrays were grown in liquid culture, the DNA was extracted with Blood & Tissue Kit (Qiagen), and the variable ends were sequenced with the Sanger method using BigDye Terminator v3.1 Cycle Sequencing Kit (Applied Biosystems) and the automated 3130xl Genetic Analyzer (Applied Biosystems).

### Population level CRISPR deep sequencing

All primer sequences are listed in Table S3. Total DNA was extracted from 1 ml liquid samples using DNeasy Blood & Tissue kit (Qiagen). The variable ends of subtype II-C and type VI-B loci (C1 and C2, respectively) from the phage and bacteria-only treatments were amplified with DreamTaq (Thermo Fisher Scientific), with one primer binding to the leader sequence (C2_B185_F and C1_B185_R) and one to the second (C1_B185_F) or third (C2_B185_R) spacer. Protocol for subtype II-C: 95°C 3 min, <u>95</u>°<u>C 30 sec, 59</u>°<u>C 30 sec, 72</u>°<u>C 1 min (cycle underlined 32x),</u> 72°C 15 min. Protocol for subtype VI-B: 95°C 3 min, <u>95</u>°<u>C 30 sec, 60.2</u>°<u>C 30 sec, 72</u>°<u>C 1 min (cycle underlined 30x)</u>, 72°C 15 min. For subtype II-C, additional magnesium was required for amplification (4 mM). To minimize PCR-bias, four separate PCR-reactions were performed for each of the five replicates and then pooled using Qiagen MinElute Reaction Cleanup kit. The resulting 10 µl samples were run on a 2% agarose gel (4.6 V/cm) for 2 h 45min in TAE buffer. Wild-type length bands (subtype II-C: 181 bp, subtype VI-B: 223 bp) and bands representing arrays with one spacer addition (subtype II-C: 246 bp, subtype VI-B: 289 bp) were extracted from the phage-treatments using X-tracta (Sigma) extraction tools and Gel Extraction Kit (Qiagen) using MinElute columns (Qiagen). In the bacteria-only treatments, the wild-type bands from both loci were extracted as controls. All extractions underwent two gel purification rounds to reduce contamination with other-length amplicons.

For deep-sequencing, we used the previously published pipeline for multiplexed Ion Torrent sequencing (*52*). Extracted type VI-B PCR-products were re-amplified using the Maxima Hot Start Taq DNA polymerase (Thermo Fisher Scientific), M13-B185_223bp_C2F and P1-B185_223bp_C2R primers, and an IonA_bc_M13 primer that contained multiplexing barcodes and an M13 adaptor (*52*). The following program was used: 95°C 3 min, 95°C 5 min, <u>94</u>°<u>C 45 sec</u>, <u>53</u>°<u>C 1 min, 72</u>°<u>C 1 min (cycle underlined 20x),</u> 72°C 5 min. As we were unable to obtain amplification of the type II-C array with the above three-primer PCR, the PCR was split in two stages. First, type II-C PCR products were amplified using the primer pair M13-B185_C1_F3 / P1-B185_181bp_C1R and the DreamTaq enzyme with added magnesium (4 mM) using the program <u>95</u>°<u>C 30 sec, 59</u>°<u>C 30 sec, 72</u>°<u>C 1 min (cycle underlined 31x),</u> 72°C 15 min. The reaction was then purified with Agencourt AMPureXP (Beckman Coulter) and re-amplified using primers IonA-bc_M13 (*52*) and P1-B185_181bp_C1R in a reaction identical to the three-primer reaction used for type VI-B. Finally, all samples were purified using the AMPureXP and quantified with Qubit fluorometer and Qubit dsDNA HS kit. Equimolar amounts (5 ng) of products were pooled for sequencing. Pooled PCR products were sequenced after emulsion PCR with the Ion OneTouch system and Ion OT2 400 kit (Life Technologies) on Ion 314 chips with Ion PGM Sequencing 400 Kit (Life Technologies), according to the manufacturer’s instructions.

### Deep sequencing data preparation

Non-trimmed reads were obtained from PGM and sorted by their barcode. Trimmomatic 0.36 (*53*) was used for quality control with the following parameters: SLIDINGWINDOW:3:21 MINLEN:100 TRAILING:23. Spacers were extracted from the trimmed reads using a custom Python script that extracted 27 to 32 bp sized spacers between an intact first repeat and the first four nucleotides of the subsequent repeat (due to short read length, a fully intact second repeat was usually not available). To obtain unique spacers, the spacer pool was clustered with CD-hit-est (*54*) using a clustering threshold of 0.8 and word size 5. Spacer sequences that were already present in the ancestral strain were identified using the FuzzyWuzzy python package and discarded.

### Mapping

The spacers were mapped to the phage genome using Bowtie 2.0 (*55*) with the following custom configuration: --very-sensitive-local --score-min G,10,8. Unmapped spacers were then mapped onto the bacterial genome and the remaining unmapped reads (most probably due to poor sequence quality) were discarded. On average, 7.9% and 1.14% of unique spacers were discarded in the type II-C and VI-B loci, respectively. The genomes were divided in bins that span 3% of the respective genome and the protospacer count of each bin was calculated in a custom Python script. The resulting protospacer distribution on both genomes was illustrated using ggplot2 package in RStudio 1.1.463 (R 3.5.3). The predicted origin of replication (oriC) region of *F. columnare* was determined using DoriC 10.0.0 (*56*).

### PAM sequences

With the aim of comparing PAM sequences between the loci, we report all PAM sequences based on adjacent DNA (instead of RNA) regions due to the large proportion of only-DNA-targeting type VI-B spacers. Similarly, we follow the guide-oriented approach in both loci, as is common with DNA-targeting systems (*57*) (PAM is depicted in the strand non-complementary to the spacer in the CRISPR transcript). All PAM sequences were extracted using custom Python scripts and shown using WebLogo (*58*).

### Spacer pool identity level analysis

To calculate the proportion of subtype VI-B spacers that are identical to II-C spacers, unique VI-B spacers from all replicates (b, d and e) were first pooled. CD-hit-est was used to extract unique spacers from the pooled spacer set (clustering threshold 0.8). The process was repeated for the type II-C spacers from replicates b and e. Next, the pooled unique spacer sets from both loci were compared to each other with CD-hit-est-2D using a similarity threshold of 0.9. The number of resulting clusters was then used to calculate the proportion of VI-B spacers that had match in the type II-C spacer pool.

Simulated sets of 430 spacers (the number of spacers in type VI-B spacer pool) were generated by sampling the phage genome either at random positions or from the 1672 predicted PAM sites (NNNNNTAAA). Sampling was repeated 1000 times and similarities of the simulated spacers was compared with the type II-C spacer pool using CD-hit-est-2D as above. A normal distribution was fit onto the simulated distributions using the R function *fitdistr* from *MASS* package. The probability of the observed similarity, given the null hypothesis, between subtypes II-C and VI-B spacers (0.58) was measured using the function *pnorm* from the upper tails of the distributions. Fig. S2 shows a graphical presentation of this result.

### Proportions of mRNA-targeting spacers

The capability of each spacer to target an ORF’s transcript was determined by two rules: (i) the crRNA of the spacer must be complementary to the coding strand and (ii) the protospacer must be fully contained within an ORF. Intergenic spacers were excluded from the analysis. The number of mRNA-targeting spacers was divided by the total number of ORF-targeting spacers in the sample to obtain the proportion of mRNA-targeting spacers for each replicate separately. We also performed the analysis for pooled spacers from the replicates. Pools were made on the basis of locus and target, resulting in four pools (II-C phage, II-C self, VI-B phage, VI-B self).

The mRNA-proportions from pooled spacers were compared to a null hypothesis that assumed an equal chance of a spacer targeting both strands. Since intergenic spacers were excluded from analysis, a binomial distribution could be used to construct the model. Separate distributions were created for each spacer pool to account for different number of spacers in each. The observed values in both loci were then compared to their respective distributions to yield direct p-values on their probability given the null hypothesis described above (one-tailed test using the *pbinom* function in R).

### RNAseq and transcription direction

Direction of CRISPR array transcription was determined from bacterial RNA-seq data. *F. columnare* B185 was grown without phage in 10 ml of Shieh medium at 24C, 150 rpm constant shaking. 24-hours grown cultures (OD 0.166-0.203) were centrifuged (5000 rpm), and stored in RNAlater (Qiagen), until RNA was extracted with Ambion MicrobExpress™ mRNA purification kit. RNA quality was verified using Agilent Bioanalyzer 2100 RNA Nano Chip and samples with RNA integrity over 9.5 selected for library preparation (Ion Total RNA-seq kit v2). The cDNA was sequenced with IonTorrent using a 318 chip (v2) and internal ERCC Spike-In control after ensuring the cDNA quality (Agilent Bioanalyzer 2100 DNA High Sensitivity chip). Our results complement previous reports, showing type VI-B crRNA transcription from the leader end (*21*) and type II-C transcription towards the leader end (starting from the repeats) or the leader-distal end of the array (*59*) (Figure S5). The RNA samples for this analysis were taken from three pooled cultures of B185 in the absence of phage, showing that these arrays are expressed constitutively, albeit at a very low level (reads mapping on arrays 167.8 and 986.6 reads per million in II-C and VI-B, respectively).

### Cas1 deletion mutant

As *F. columnare* B185 was unable to receive plasmids via conjugation or electroporation, we used *F. columnare* B245 to create a *cas1*-deletion mutant (Δcas1). The 2.1 kbp region upstream of *cas1* was amplified by PCR using Phusion DNA polymerase (New England Biolabs) and primers 2322 (adding a KpnI site) and 2323 (adding a BamHI site). The product was digested with KpnI and BamHI and ligated into pMS75 (*60*) that had been digested with the same enzymes, to produce pRC30. A 495 bp region downstream of *cas1* was amplified using primers 2324 (adding a BamHI site) and 2325 (adding a SalI site). The product was digested with BamHI and SalI and ligated into pRC30 that had been digested with the same enzymes, to generate pRC32. The plasmid pRC32 was transferred from *E. coli* S17-1λpir into *F. columnare* strain B245 by conjugation. One ml of *E. coli* overnight culture was inoculated into nine ml fresh LB containing 100 µg/ml ampicillin and incubated with shaking at 37°C until the OD_600_ reached 0.6. Similarly, five ml of *F. columnare* overnight culture was inoculated into 25 ml fresh TYES broth (*61*) and incubated at 28°C with shaking, until the OD_600_ reached 0.6. *E. coli* and *F. columnare* cells were centrifuged at 5000 rpm for 15 minutes, the pellets were each washed with 10 ml of TYES medium and centrifuged at 5000 rpm for 10 minutes. The *E. coli* and *F. columnare* cell pellets were each suspended in 0.8 ml of TYES, mixed together, and centrifuged at 7000 rpm for 3 minutes. Excess media was removed, and the mixed pellet was suspended and spotted on FCGM agar medium (*62*). After incubation at 30°C for 24 hours, cells were scraped off the plate and suspended in 1.5 ml of TYES medium. 100

µl aliquots were spread on TYES agar containing 1 µg/ml tobramycin and 5 µg/ml tetracycline and incubated at 30°C for 72 hours. The resulting tetracycline-resistant colonies were streaked for isolation, inoculated into TYES media without tetracycline and incubated overnight at 25°C with shaking to allow plasmid loss. The cells were plated on TYES media containing 10% sucrose, incubated at room temperature (20°C) to select for lack of sucrose toxicity, and were streaked for isolation using the same selection. Positive deletions were screened with PCR and gel electrophoresis, and verified with Sanger sequencing using primers Cas1_F and Cas1_R. All primers used are listed in Table S3.

Strains used in the cas1-deletion mutant spacer acquisition experiment were Δcas1 and B245_rev (rev-wt). Rev-wt is a reversion mutant where the integrated plasmid was lost by recombination in a manner that regenerated the wild-type sequence, instead of the *cas1*-deletion. The lytic dsDNA phage V156 (*18*) was used in the spacer acquisition experiment. Infectivity of V156 against the two hosts was measured with a standard double-layer method, where 300 µl of a turbid bacterial culture was mixed with 3 ml of TYES with 0.1% mucin and 1% agar. The mixture was poured on a TYES agar plate and 10 µl of phage V156 dilutions (10^−1^ to 10^−7^) were spotted in duplicates on the solidified agar. Number of plaques was counted after two days from the 10^−7^ dilutions. Phage titers (pfu/ml) were 1.9×10^10^ and 3.7×10^10^ in Δ*cas1* and rev-wt, respectively.

For the spacer acquisition experiment, both strains were grown overnight in single 5ml cultures to reach OD 0.45. Ten phage-containing and two bacteria-only 0.1X TYES cultures were started from these cultures. The spacer acquisition protocol differed from the one used with B185 as follows: medium TYES (vs. SHIEH), volume 1 ml (vs. 5 ml), duration of growth in 0.1X medium three weeks (vs. five days). After three weeks in diluted medium (0.1X), the cultures were transferred to regular TYES medium (1X) and new spacers sampled after two days. All 12 cultures of Δ*cas1* showed growth in the 1X medium in contrast to only two replicates (both containing phage) of the rev-wt. The variable ends of both arrays in all surviving cultures were amplified using primers B245_C1_F + F_col_C1_R (B245 II-C array) and F_col_C2_F + B245_C2_R (B245 VI-B array) (Figure S6).

### Comparison of type II-C and VI-B leaders and repeats in other species

Species that carry intact subtype II-C (containing *cas1*, *cas2*, *cas9* and a CRISPR array) and VI-B (containing *cas13b* and a CRISPR array) loci were listed using CRISPRCasdb (*63*) and one strain per species was selected for further analysis (*Flavobacterium branchiophilum* was excluded from analysis due to the absence of *cas1* in the subtype II-C locus). From each strain, repeat sequences from both loci were aligned with Geneious aligner (global alignment with free gaps, 51% cost matrix, gap open penalty 12, gap extension penalty 3). 200 bp leader sequences were extracted downstream of the expected variable end of the array (determined by the repeats’ direction as compared to *F. columnare* B185). To allow for meaningful comparison of the highly divergent leaders, the strains were first clustered based on their Cas1 protein (Jukes-Cantor, UPGMA). The leaders from both loci within the resulting six clusters (most of which contained single species) were aligned with Geneious aligner (global alignment, 65% cost matrix, gap open penalty 17, gap extension penalty 12). We used strict alignment rules to emphasize the importance of distance of possible motives from the repeat-leader junction. Analyses and alignments were performed with Geneious 9.1.8 (https://www.geneious.com).

