## Supplementary File for "Cooperation between CRISPR-Cas types enables adaptation in an RNA-targeting system"

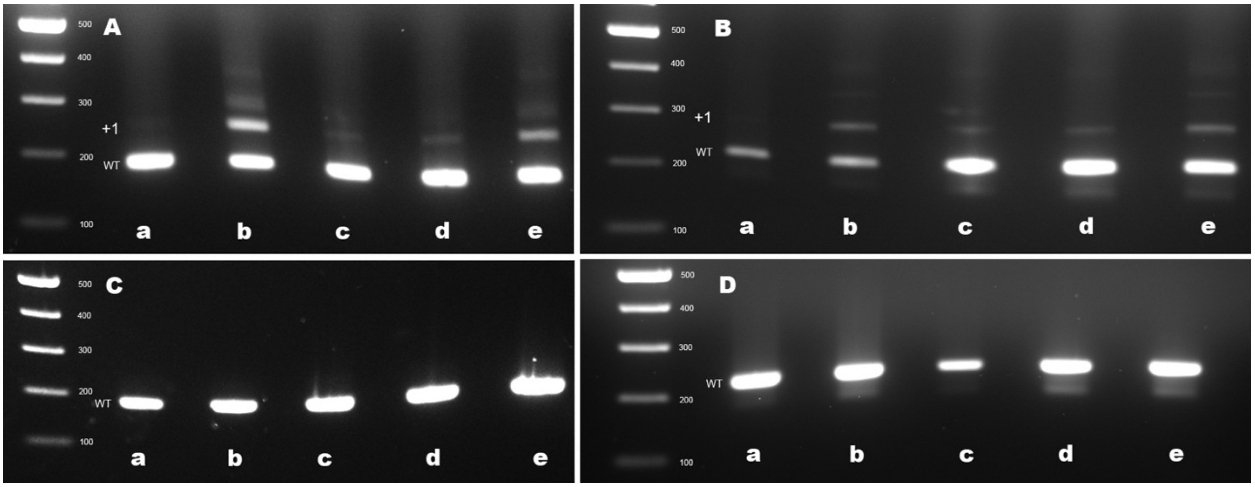


**Fig. S1.**
Gel electrophoresis of type II-C and VI-B CRISPR arrays in *Flavobacterium columnare* B185. The variable ends of the arrays were amplified from population-level DNA samples of five replicate cultures (a-e) in the presence or absence of phage FCL2. A) II-C array with phage, B) VI-B array with phage, C) II-C array without phage D) VI-B array without phage.

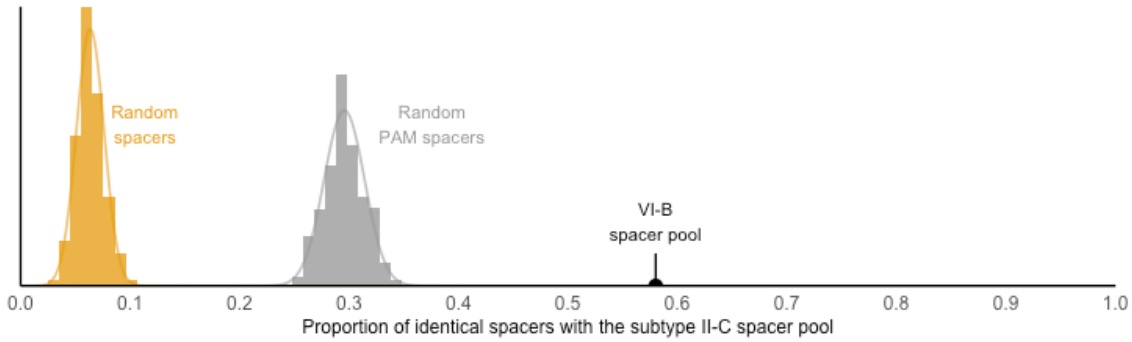


**Fig. S2.**

The proportion of common spacers between simulated or VI-B spacers with subtype II-C spacers. Compared to the pooled II-C spacers, randomly sampled spacers have a mean similarity of 6.3% (SD 1.2%) and PAM-adjacent spacers a mean similarity of 29.6% (SD 1.85%). More than half (58%) of subtype VI-B spacers are found in the II-C pool. The number of spacers in the simulated pools equal the number of spacers in the subtype VI-B pool.

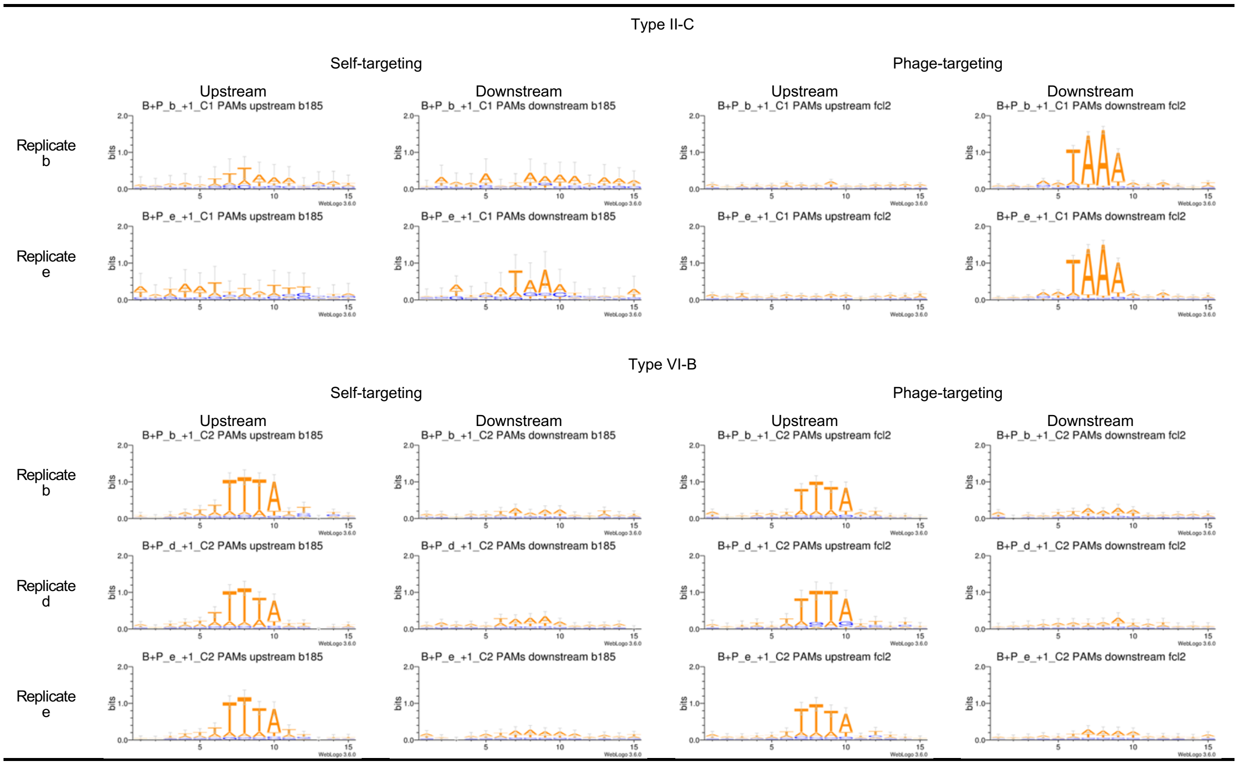


**Fig. S3.**

PAM sequences. 15 bp regions upstream and downstream of each protospacer displayed as Weblogo from replicates b and e for type II-C and replicates b, d and e for type VI-B. PAMs are determined using the guide-oriented approach.

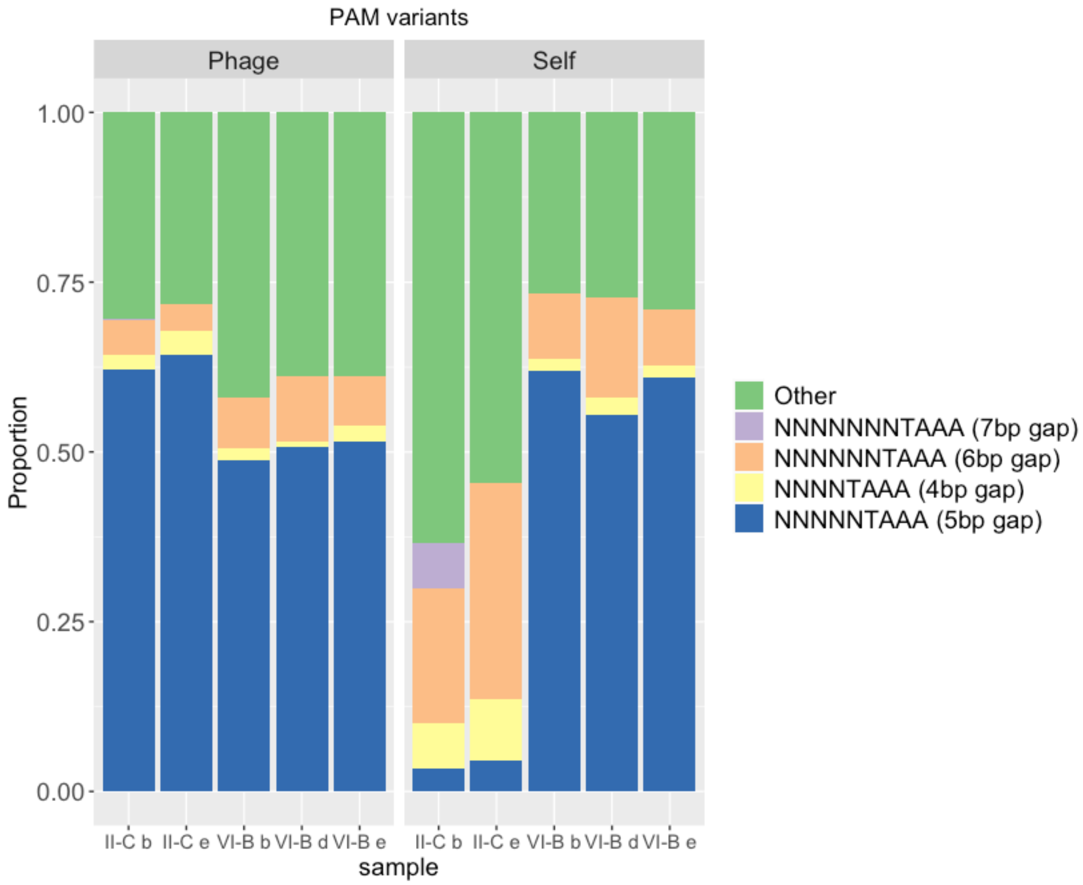


**Fig. S4.**

Variants of PAMs in both loci. Stacked bars show the proportions of different variants of PAMs in both loci and target genomes. Type VI-B PAMs have been transposed from the target-specific reverse-complement sequence (TTTANNNNN) to simplify the plot.

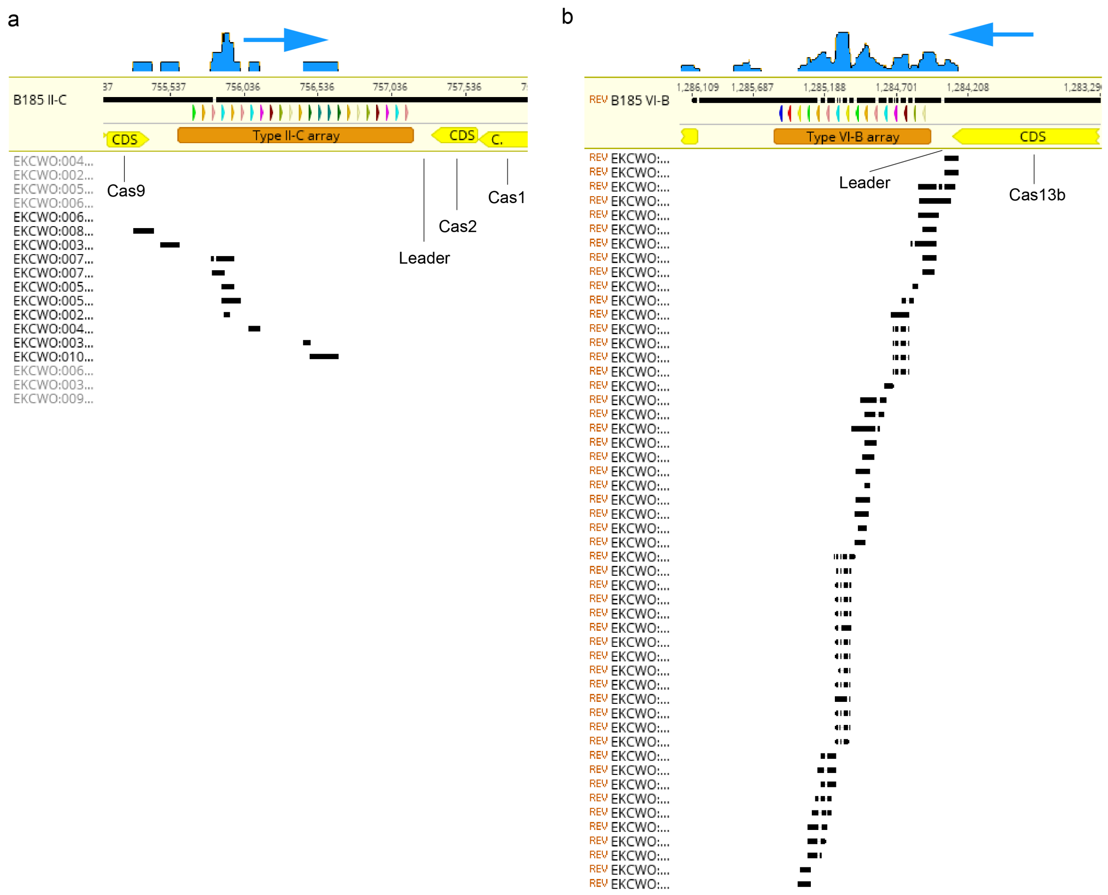


**Fig. S5.**

Transcription directions of type II-C and VI-B arrays. The RNA samples were taken in the absence of phage to provide enough RNA for sequencing, leading to a minimal yet sufficient expression of the arrays to determine transcription direction. A) 8 reads revealed type II-C transcription from within the array towards the leader (167.8 reads per million). b) 47 reads mapped on the type VI-B array revealed transcription direction starting from the leader end (986.6 reads per million).

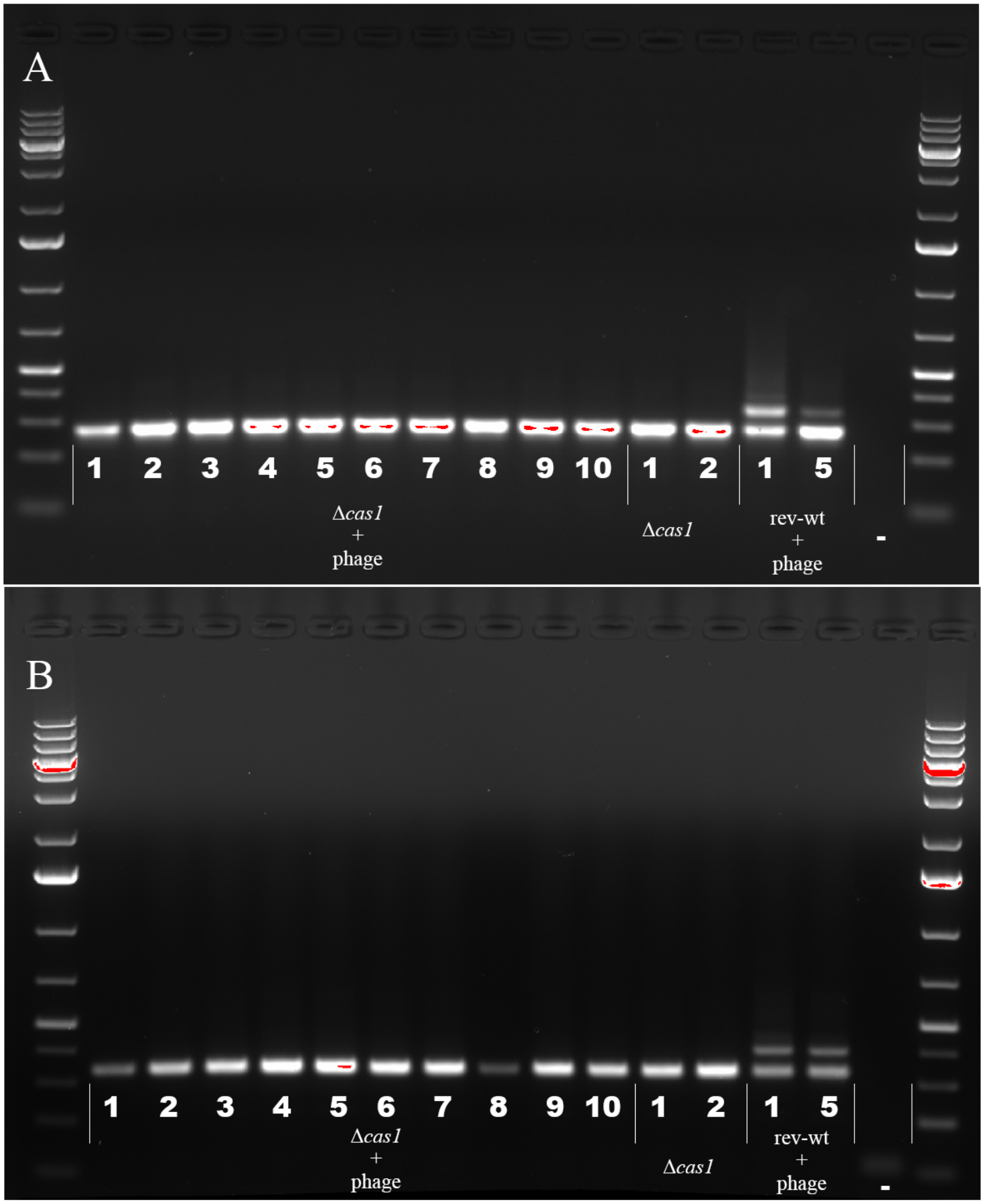


**Fig. S6.**

Spacer acquisition experiment using *Δcas1*. A) Subtype II-C array B) Subtype VI-B array. Ten replicates of *Δcas1* and rev-wt (annotated with numbers) were grown with phage along with two replicates without phage. Of the 12 rev-wt cultures, only two had grown at the end of the three-week experiment, both having acquired new spacers in the II-C and VI-B CRISPR-Cas loci. All *Δcas1* replicates were grown at the end of the experiment, but none had obtained new spacers in either locus. Ladder used on gels: 1 kb Plus DNA Ladder (Thermo Fisher Scientific).

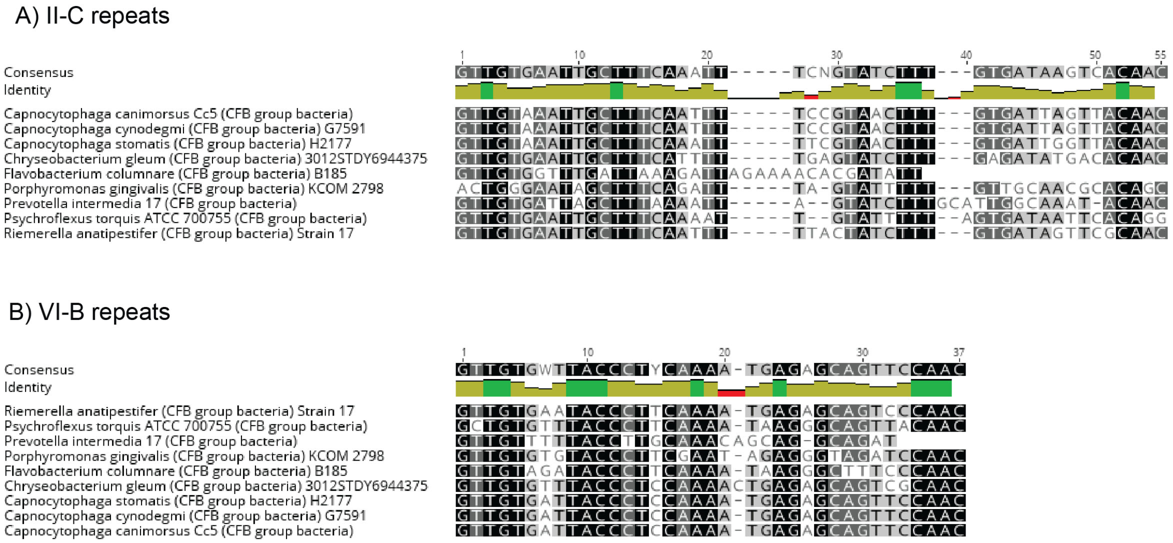


**Fig. S7.**

Alignments of II-C and VI-B repeats from species that carry both loci. 3’ ends are leader-adjacent.

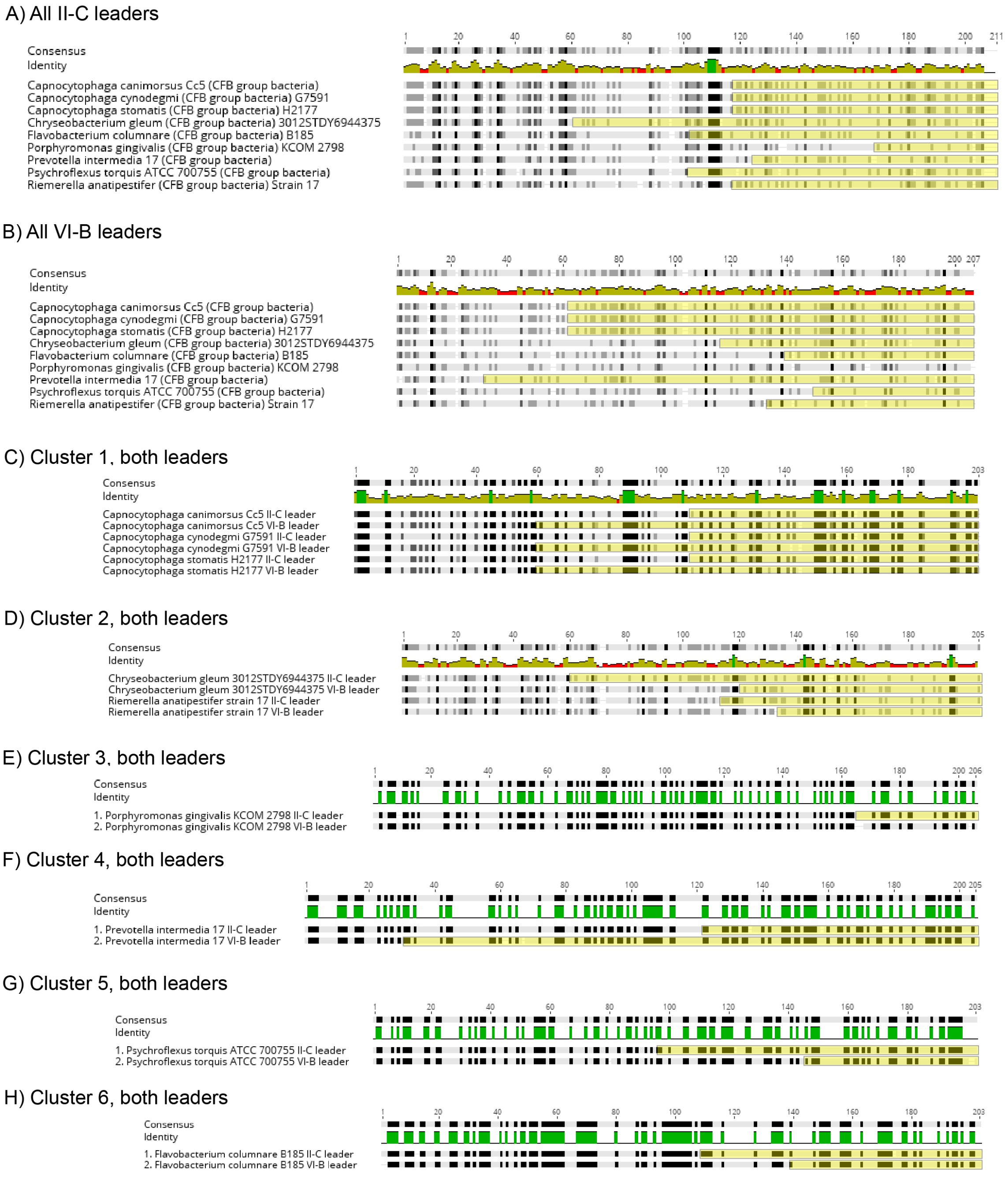


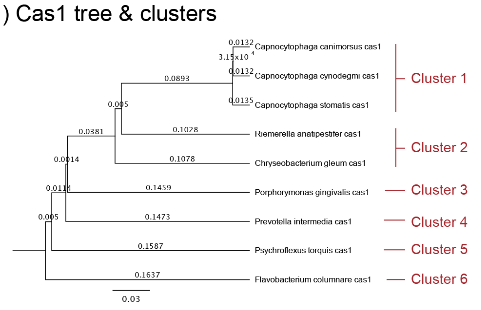


J) Similarity of leaders in clusters

| Cluster | II-C & VI-B leader similarity |
|---|---|
| *1. Capnocytophaga (canimorsus, cynodegmi, stomatis)* | 58.6 % |
| *2. Chryseobacterium gleum, Riemerella anatipestifer* | 38.7 % |
| *3. Porphyromonas gingivalis* | 40.3 % |
| *4. Prevotella intermedia* | 42.3 % |
| *5. Psychroflexus torquis* | 41.9 % |
| *6. Flavobacterium columnare* | 47.8 % |

**Fig. S8.**

Alignments of II-C and VI-B leaders from species that carry both loci. 5' end is repeat-adjacent and open reading frames downstream of the leader are annotated with yellow rectangles. A) Alignment of type II-C leaders (200 bp). B) Alignment of type VI-B leaders (200 bp). C-H) Alignments of both leaders within clusters. I) Phylogenetic tree of Cas1 and the derived clusters. J) Nucleotide similarity of leaders in clusters.

**Table S1.**

Colonies with additional spacers and their sequences. ORF numbers indicate phage genes and "self" indicates bacterial chromosomal genes.

| Mutant ID | Locus | Colony morphotype | Spacer target | Spacer sequence |
|---|---|---|---|---|
| 1c30z | II-C | Rhizoid | FCL2 (ORF64) | AAGTGCTAACTTCAATACGCTGCATGGTGG |
| 1a5z | VI-B | Rhizoid | Self (RHS repeat-associated core domain protein) | CAACTTGTCATTCTCATCAAAAGGCCCTTC |
| 1c13z | VI-B | Rhizoid | Self (diacylglyceryl transferase) | TAAGATGTCCTTTTTAATTATTTTTCAGAG |
| 1c38r | VI-B | Rough | FCL2 (ORF58) | AACGATTTAGCGTTAATAATTTCGTTCGA |
| 1d14z | VI-B | Rhizoid | FCL2 (ORF77) (1bp over ORF) | TTCTGCCCTCTAAATTTAATATCTCTCATA |
| 1c5z | 1x II-C, 2x VI-B | Rhizoid | II-C: FCL2 (ORF58)<br>VI-B: FCL2 (ORF64) & self (damage-inducible protein) | AAATTTGTTTACAATTTGAACTTGTTCCAG,<br>TAAAATTTTATTGTTTTTAAACCGCTTTC,<br>TTACCGGGAGTGCCTTATGAAATGAAACAC |

**Table S2.**
Numbers of individual colonies screened and the resulting number of CRISPR mutants

| CONDITION | REPLICATE | SCREENED | NUMBER OF MUTANTS IN INITIAL SCREEN | NUMBER OF MUTANTS AFTER SERIAL PLATING |
|---|---|---|---|---|
| B + P + UVP | A | 15 | 5 | 1 |
| B + P + UVP | B | 3 | 0 | 0 |
| B + P + UVP | C | 40 | 8 | 4 |
| B + P + UVP | D | 14 | 1 | 1 |
| B + P + UVP | E | 13 | 0 | 0 |
| B+ P | A | 20 | 0 | 0 |
| B+ P | B | 0 | 0 | 0 |
| B+ P | C | 20 | 0 | 0 |
| B+ P | D | 21 | 0 | 0 |
| B+ P | E | 19 | 0 | 0 |
| B + UVP | A | 20 | 0 | 0 |
| B + UVP | B | 20 | 0 | 0 |
| B + UVP | C | 7 | 0 | 0 |
| B + UVP | D | 10 | 0 | 0 |
| B + UVP | E | 20 | 0 | 0 |
| B | A | 4 | 0 | 0 |
| B | B | 4 | 0 | 0 |
| B | C | 7 | 0 | 0 |
| B | D | 7 | 0 | 0 |
| B | E | 17 | 0 | 0 |

**Table S3.**
Primers used in the study

| Primer | Info | Sequence |
|---|---|---|
| C2_B185_F | Type VI-B array, B185 specific | ACTATGAGAGGCTTACCAGCGTTTA |
| C2_B185_R | Type VI-B array, B185 specific | CAAACACATTGTATCATTTAGCAGTAA |
| C1_B185_F | Type II-C array, B185 specific | TGTTCTATCGGCGTAAAATAAGTT |
| C1_B185_R | Type II-C array, B185 specific | TAACGATATTTCGGCTTAATGCT |
| M13-B185_223bp_C2F | Type VI-B, B185, M13 underlined | TGTAAAACGACGGCCAGTACTATGAGAGGCTTACCAGCGTTTA |
| P1-B185_223bp_C2R | Type VI-B, B185, P1 underlined | CCTCTCTATGGGCAGTCGGTGATCAAACACATTGTATCATTTAGCAGTAA |
| P1-B185_181bp_C1R | Type II-C, B185, P1 underlined | CCTCTCTATGGGCAGTCGGTGATTAACGATATTTCGGCTTAATGCT |
| M13-B185_C1_F3 | Type II-C, B185, M13 underlined | TGTAAAACGACGGCCAGTTGTTCTATCGGCGTAAAATAAGTT |
| B245_C1_F | Type II-C, B245 specific | CTGTTTTGTTTCATTTGGTAAATCA |
| B245_C2_R | Type VI-B, B245 specific | GATGTAGAAATACTTAGCGACAATGTAG |
| F_col_C2_F | Type VI-B, F. columnare common | GGTCTAAATACAATTGCTCTTTGACATT |
| F_col_C1_R | Type II-C, F. columnare common | CCCTAAAGCACCACAACCCA |
| B245_cas1_F | Cas1 sequencing primer (B245) | TGTACATCGGCACTTTCGCT |
| B245_cas1_R | Cas1 sequencing primer (B245) | AGTATTCCCGCCCCGTATTT |
| 2322 | B245 cas1 upstream fragment F | GCTAGGGTACCAACGGATGGAGCAATAAGTGT |
| 2323 | B245 cas1 upstream fragment R | GCTAGGGATCCAGTAGCCTCTCGTAAAATCCC |
| 2324 | B245 cas1 downstream fragment F | GCTAGGGATCCGCAAGTTCCTTACAACAGTGT |
| 2325 | B245 cas1 downstream fragment R | GCTAGGTCGACAGCTAACAGCTTAGTTATTAAATATCAAAG |
| 2367 | Plasmid sequencing primer | GCTAGGGTACCAATTCGATACCCGTTTAAATCCA |
| 2368 | Plasmid sequencing primer | GCTAGGGATCCCTAAAAGAGCTGTTATATTCATCG |
